# Adaptive reprogramming during early seed germination requires temporarily enhanced fermentation – a critical role for alternative oxidase (AOX) regulation that concerns also microbiota effectiveness

**DOI:** 10.1101/2021.06.08.447570

**Authors:** Bharadwaj Revuru, Carlos Noceda, Mohanapriya Gunasekaran, Sarma Rajeev Kumar, Karine Leitão Lima Thiers, José Hélio Costa, Elisete Santos Macedo, Aprajita Kumari, Kapuganti Jagadis Gupta, Shivani Srivastava, Alok Adholeya, Manuela Oliveira, Isabel Velada, Debabrata Sircar, Ramalingam Sathishkumar, Birgit Arnholdt-Schmitt

## Abstract

Plants respond to environmental cues via adaptive cell reprogramming that can affect whole plant and ecosystem functionality. Microbiota constitutes part of plant’s inner and outer environment. This *Umwelt* underlies steady dynamics, due to complex local and global biotic and abiotic changes. Hence, adaptive plant holobiont responses are crucial for continuous metabolic adjustment at systems levels. Plants require oxygen-dependent respiration for energy-dependent adaptive morphology, such as, germination, root and shoot growth, formation of adventitious, clonal and reproductive organs, fruits and seeds. Fermentative paths can help in acclimation and, to our view the role of alternative oxidase (AOX) in coordinating complex metabolic and physiologic adjustments is underestimated.

Cellular level of sucrose is an important sensor of environmental stress. We explored the role of exogenous sucrose and its interplay with AOX during early seed germination. We found that sucrose-dependent initiation of fermentation during the first 12 hours after imbibition (HAI) was beneficial to germination. However, parallel enhanced AOX expression was essential to control negative effects by prolonged sucrose treatment. Early down-regulated AOX activity until 12 HAI improved germination efficiency in the absence of sucrose, but suppressed early germination in its presence. Our results also suggest that seeds-inoculated arbuscular mycorrhizal fungi can buffer sucrose stress during germination to restore normal respiration more efficiently.

Following this approach, we propose a simple method to identify organic seeds and low-cost *on-farm* perspectives for early selection on disease tolerance, predicting plant holobiont behavior and improving germination. Furthermore, our research strengthens the view that AOX can serve as powerful functional marker source for seed hologenomes.

## Introduction

Understanding the role of microbiota in adaptive plant robustness is important for crop improvement and for developing innovative tools that could allow more efficient plant selection (Arnholdt-Schmitt et al., 2014, Nogales et al., 2015, Arnholdt-Schmitt et al., 2015, Arnholdt-Schmitt et al., 2018). Research on the relevance of endophytic and associated microbiota and usage of microbiota inoculation are often hampered by low reproducibility, which is due to a lack of better understanding the fundamental principles of functional plant-microbiota interaction (Arnholdt-Schmitt, 2008, Vicente and Arnholdt-Schmitt, 2008, Mercy et al., 2015, Campos et al., 2015, Mercy et al., 2017, Bedini et al., 2018). Albornoz et al. (2020) emphasizes the need for studying mycorrhizal benefits on a case-by-case basis that should consider more holistic and context-dependent views on mycorrhiza functioning at plant family- and biome-wide levels. Also, it is widely confirmed that endophyte effects are genotype-specific (Abdelrazek et al., 2020a,b). Further, Durán et al. (2018) identified bacterial endophytes as drivers for soil suppressive take-all disease. Nevertheless, they highlighted that they did not find relevant correlation between disease suppression and reduced pathogen biomass. In our opinion, these are key observations. They encourage us to continue working on the hypothesis that individual plant host’s competence for resilience plays the most critical role for beneficial or non-beneficial plant – microbiota interaction, which can be superior to plant families and biome origins.

However, there is lack of knowledge on traits that aid in (a) early prediction of the strength of plants and (b) demonstration of its relevance for plant-microbiota interactions and (c) transformation of such knowledge into user- and environment-friendly applications for sustainable agriculture. We earnestly aim with our perspective to understand these phenomena and to contribute to the knowledge base towards closing these three gaps.

The capacity for efficient reprogramming as a trait *per se* is recognized as marker for adaptive plant robustness (Cardoso and Arnholdt-Schmitt, 2013). Seed germination can serve as an experimental *in vitro* tool to study environmental stress-induced reprogramming and to identify early functional markers and tools for predicting plant performance under field conditions (Mohanapriya et al., 2019). Dry seeds are known to respond to water imbibition and subsequent penetration of oxygen. Thus, radicle emergence can be seen as an indicator of environmental stress recovery from the dry-to-water imbibed conditions and low-to-high oxygen status.

Efficient seed germination under field conditions is especially required in organic agriculture, where application of chemical herbicides for suppressing weed competition and pesticides shall be avoided in support of healthy food and feed production and to improve sustainability of bio-based socio-economic systems. At the same time, organic agriculture impacts seeds quality and the amount of microbiota in seeds (Naomi Cope-Selby et al., 2017, Wassermann et al., 2019). Recently, the use of the so-called ‘organic seeds’ versus conventionally produced seeds is raised as an ethical issue (www.liveseed.eu; effective European regulation from 1.1.2021: EC No 2018/848). However, the better quality of organic seeds in terms of their contribution to agriculture sustainability, nutritional quality and yield performance is under intensive debate (e.g. Voss-Fels et al., 2019, Bhaskar et al., 2019) and requires scientific clarification (Simon et al., 2017, Abdelrazek et al., 2020a,b). Appropriate methods and tools are the need of the time in order to discriminate organic versus conventional seeds by traits that should at the same time allow predicting the assumed superior quality of organic seeds.

## Background

Cellular reprogramming is an energy intensive phenomenon. Reactive oxygen species (ROS) are known to interact with redox-sensitive protein cysteine thiol groups relevant for energy metabolism and metabolic channeling linked to cell differentiation and cell cycle regulation (Bigarella et al. 2014, Dumont and Rivoal, 2019; Qi et al., 2020; Gupta et al. 2020a, Gupta et al., 2020b, Pengpeng et al., 2020). Sugars and sugar phosphates interact with hormone-mediated signaling networks to modulate energy metabolism. Auxin-stimulated sugar metabolism was frequently reported (e.g. Zhao et al., 2021). But only few examples revealed that sucrose can induce new cell programs (Grieb et al., 1994, see in Zavattieri et al., 2010) and also, vice versa, can change auxin metabolism (Lin et al., 2016, Meitzel et al., 2020). In maize, sucrose stimulated more cell cycle markers during germination than glucose (Lara-Núñez et al., 2017). Down-stream of sugars, two important antagonistic protein kinases are involved in energy sensing and physiological adaptation (reviews in Bayey-Serres et al., 2018, Schmidt et al., 2018, Sakr et al., 2018). While sucrose non-fermenting-1-related protein kinase1 (SNRK1) is activated when energy is depleted (Schmidt et al., 2018, Wurzinger et al., 2018; Wang et al., 2020), TOR (target of rapamycin) is induced under conditions of energy excess and stimulates cell cycle progression and cell proliferation (Sangüesa et al., 2019). Sucrose can have various functions: besides its nutritional role it acts as signaling component (Baena-Gonzalez et al., 2017, Sakr et al., 2018), as osmotic stressor that can disrupt communication within and between cells (Moon et al., 2015) and was shown to trigger aerobic alcohol fermentation in support of respiration and biosynthesis of higher molecular weight compounds, such as lipids (Mellema et al., 2002). Alcohol fermentation was found to play critical role in controlling tissue level pyruvate in plants and, thereby, adapt respiration rates to the prevailing cellular energy status (Zabalza et al., 2009). Fan et al. (2020) identified hormone and alcohol degradation pathways as the most activated during early stages of somatic embryogenesis (SE), which is a prominent example of *de novo* programming. Ethanol has been shown to reduce ROS levels and led to high induction of alternative oxidase (AOX) and glutathione-S-transferase transcripts (Nguyen et al., 2017). Transcriptome analyses at 2,4-D - induced reprogramming indicated that the extent of aerobic fermentation is connected to cell proliferation and was regulated by interacting levels of sucrose and AOX (Costa et al., 2021; preprint). Transient up-regulation of genes related to alcoholic and lactic fermentation was shown to be associated with glycolysis and modified complex stress signaling patterns with enhanced superoxide dismutase and decreased transcript levels of nitric oxide - producing nitrate reductase. Further, our data signaled activation of cell death-regulating system and arrested cell cycles at reduced alpha-tubulin transcription at the earliest step in reprogramming. Considering generality of these observations, we proposed a reference transcriptome profile to identify virus traits that link to harmful reprogramming (Arnholdt-Schmitt et al., 2021 In Press). This approach helped identifying an early trait for combating SARS-CoV-2 that covers ROS/RNS balancing, aerobic fermentation regulation and cell cycle control (Costa et al., 2021; preprint).

In seeds, fermentation and alternative respiration are dominating (references in Arnholdt-Schmitt et al., 2018, Mohanapriya et al., 2019). During seed germination, structural and functional acclimation of aerobic respiration is central and determines temperature-dependent efficiency of germination (Bello and Bradford, 2016, Gaël Paszkiewicz et al., 2017). Nevertheless, markers for respiration and oxygen consumption were not superior to simple germination tests for predicting seed vigor from single seeds (Powell et al., 2017). However, it was suggested that alternative respiration plays the most critical role during germination (Arnholdt-Schmitt et al., 2018 and references herein, Mohanapriya et al., 2019). This role requires managing ROS/RNS increase and channeling energy and substance flow from fermentation when carbohydrate storages are released and enzymes get into motion (Saleh and Kalodimos, 2017), but the respiration chain is still structurally restricted and overloaded at massively incoming oxygen. AOX is mainly regulated by pyruvate (Millar et al, 1996, Hoefnagel et al., 1997, Albury et al., 2009, Hakkaart et al., 2006, Carré et al., 2011, Selinski et al., 2018) and, strikingly, Ito et al. (2020) showed in *Arum* that energy-related metabolic regulation can be determined by temperature-dependent switching between AOX polymorphisms in the binding site for AOX-pyruvate. In this scenario, it might be of interest that AOX was essential in ethylene-induced drought tolerance and mediating autophagy generation via balancing ROS levels (Zhu et al., 2018). Thermo-inhibition of carrot seed germination could be circumvented by seed priming, which was found to be linked to increased ethylene production at higher temperatures (Nascimento et al., 2013). Ethylene biosynthesis was found to be induced by H2O2 and acted positively on germination independently from auxin-coordinated hormonal crosstalk linked to ABA suppression and gibberellin activation (Wojtyla et al., 2016). During plant ethylene biosynthesis cyanide is generated as by-product of the pathway and suspected to help shifting cyt respiration to alternative respiration (Siegieñ and Renata Bogatek, 2006, Machingura et al., 2016). Eckert et al. (2014) stressed that microbiota have developed ethylene-producing pathways to profit during invasion and to evade from defense responses of the host plants. Mercy et al. (2017) observed that KCN treatment of mycorrhizal seedlings promoted local arbuscular formation.

Recently, we identified AOX as stress level - sensing coordinator for auxin-inducible metabolic reprogramming by comparing induction of SE and seed germination (Mohanapriya et al., 2019; see also Arnholdt-Schmitt et al., 2018). Association of AOX to target cell reprogramming was also observed in other systems such as adventitious root induction in olive (Macedo et al., 2009, Porfirio et al., 2016) and elicitor-induced hairy roots (Sircar et al., 2012). Furthermore, our group had contributed to novel functional marker strategies by highlighting AOX as a marker across taxonomic boarders that considered ‘shared’ *AOX* genes in plant holobionts (Arnholdt-Schmitt 2005a and b, Arnholdt-Schmitt et al., 2006, Arnholdt-Schmitt, 2008, Campos et al., 2015, Mercy et al., 2017, Bedini et al., 2018). Based on the role of AOX in carbohydrate metabolism (Vanlerberghe et al., 1994), our approach stimulated reflecting on the role of fermentation and sugars for plant –mycorrhiza interaction (Mercy et al., 2017, Bedini et al., 2018) and had led to a privately explored patent (Mercy and Mercy, 2014). However, the early phase of reprogramming was not sufficiently considered in that research (Mercy et al., 2017) to drive our core functional marker approach (Arnholdt-Schmitt, 2008, Mercy et al., 2015). Recently, Mohanapriya et al. (2019) observed that AMF inoculation in imbibed seeds interacted with the AOX-inhibitor SHAM and palliated negative SHAM effects on early germination. Also, AMF effects in seeds seemed to be modified by non-culturable microbiota. Integrated *in silico* studies on experimental data revealed that endophytes interact with AOX expression in a species-, stress-, and developmental-dependent manner. *Enterobacter* species could reduce salt-induced expression of AOX1a and kept its mRNA level low when applied together with salt. Costa et al. (2021; preprint) highlighted that microbiota - plant genotype interaction and its impact on early carrot seed germination can be modified by SHAM.

In our earlier work (Mohanapriya et al. 2019), we demonstrated successful prediction by oxycaloric equivalents from germinating seeds at 10 HAI. The present perspective questions the metabolic nature of AOX coordination and provides deeper phenotyping during germination of endophyte-free and microbiota-inoculated seeds focused at early times around 12 HAI. **Figure 1** demonstrates the step-by-step rationale of fundamental insights and deduced practical strategies.

**Figure 1:**
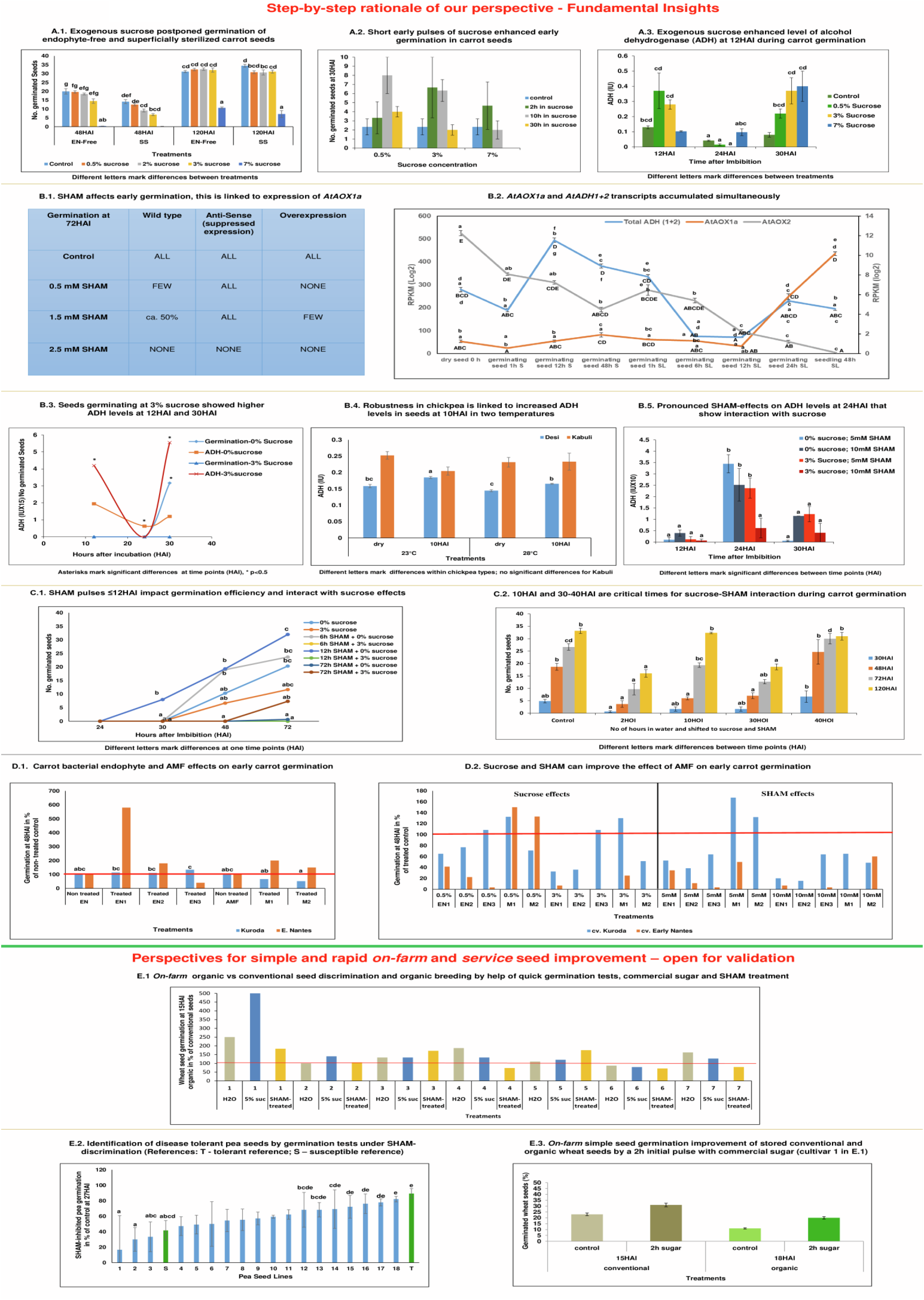
Step-by-step rationale of our perspective. **A.1 Exogenous sucrose postponed germination of endophyte-free (EFS) and superficially sterilized (SSS) carrot seeds**: sucrose inhibited early germination (at 48 hours after imbibition (HAI)) dependent on increasing sugar concentrations. This trend was the same for seeds treated to become endophyte-free and seeds that were superficially sterilized. At 120 HAI, the effect of 0.5 to 3% sucrose could not be noticed anymore, while 7% sucrose inhibited germination for a prolonged time. This observation indicates a critical role of sucrose during induction of adaptive performance. For confirmation of this role of sucrose, in supplementary **Figure S1**, the effect of sucrose is shown for auxin-dependent early induction of somatic embryogenesis (SE) as the most studied example of *de novo* programming. It demonstrates that (a) sugar is essential for cell reprogramming, since SE induction was not observed at around 45 DAI in controls, but only at 2% and 3% sucrose supply and (b) that SE can be optimized with the help of increasing amounts of exogenous sucrose, since SE induction efficiency was highest at 3% sucrose (Supplementary table S1). Cell reprogramming competes with cell division. This is a common insight, which got here validated again through the observed delay in embryonic versus non-embryonic callus emergence by increasing sucrose concentrations at lower levels. As a general tendency, at increasing sucrose levels, less seeds showed callus growth, which later demonstrated to be embryogenic, in comparison to the higher number of seeds with (non-embryogenic) callus growth at low sucrose levels (Figure S1). **A.2 Short early pulses of sucrose enhanced early germination in carrot seeds:** 3% sucrose applied for 2 h or 10 h from imbibition enhanced early germination to about the same degree compared to the control and to a longer pulse of 30 h. A lower sucrose concentration of 0.5% had the highest effect only by a longer pulse of 10 h and, at 7% sucrose a higher effect against the control was only indicated when given as a short pulse of 2 h. This observation was confirmed with a second carrot cultivar in a rapid *on-farm* check by using a ca. 5% solution of commercial sugar (significant) (**Figure S2**). **A.3 Exogenous sucrose enhanced the level of alcohol-dehydrogenase (ADH) at 12HAI during carrot germination:** At 12 HAI, treatment with 0.5% and 3% sucrose resulted in an increase in ADH activity as compared to the control, while for 7% sucrose no effect was observed. At 24 HAI, the control indicated decline of ADH values. In the presence of 0.5% and 3% sucrose, this decline was not avoided or might even have been strengthened. However, at 30 HAI, a second phase started, where sucrose enhanced the level of ADH in a concentration-dependent manner including a positive effect of 7% sucrose. **B.1 SHAM affects early germination and this links to expression of *AtAOX1a***: In wildtype *Arabidopsis thaliana* seeds, monitoring germination at 72 HAI showed that SHAM treatment led to reduced germination rates. This inhibition was dependent on its concentration of 0.5 or 1.5 mM. However, when AOX had been silenced (AS), SHAM did not affect germination. To the contrary, when AOX was constitutively overexpressed (OE), SHAM indicated stronger inhibition of germination than in the wildtype. Nevertheless, the three genotypes germinated with similar efficiency in relation to their respective controls. This latter observation points to the fact that AOX is critically important for germination, if present. However, in case it is not present or activated (AS) alternative pathways can substitute the functional role of AOX during germination. **B.2 *AtAOX1a* and *AtADH1+2* transcripts accumulated simultaneously**: a study on ADH transcript accumulation in wildtype *Arabidopsis thaliana* confirmed a biphasic activation of *ADH* during germination. A first increase was observed 12 h after stratification (significant), which includes imbibition of water. The second enhancement occurred from 12 h SL shortly before root emergence was monitored at 24h SL. In parallel to increased *ADH* transcript accumulation, *AOX1a* transcripts accumulated during both phases, i.e. induction and early initiation of germination. After early induction, *ADH* transcripts showed a high decline (significant) until the end of the dark stratification phase, while *AOX1a* transcript levels remained more stable. During the second phase at initiation of exponential root length growth in light observed at 48h SL, *AOX1a* transcript accumulation keeps on enhancing, while the increase of ADH transcripts stopped at that time point. This was also indicated at the first phase. *AOX2* transcript accumulation was differentially regulated in comparison to *AOX1a* and showed continuous down-regulation during the whole period, which appeared to be stronger in the SL phase. **B.3 Seeds germinating at 3% sucrose showed higher ADH levels at 12HAI and 30HAI:** during early germination of carrot seeds, ADH levels follow a parable, when monitored between 12 and 30 HAI. This was observed in control seeds and seeds germinating at 3% sucrose. Nevertheless, suppressed germination at 3% sucrose was linked to higher levels of ADH at 12 HAI and at 30 HAI. This means, the more efficient germination in control seeds was linked at these two time points to lower levels of ADH. Under both conditions, in the absence of exogenous sucrose and at 3% sucrose, 24 HAI displayed a turning point with lowest ADH activity levels. However, ADH activity at 24 HAI was higher in controls (significant) than under conditions of sucrose-supplementation. **B.4 Robustness in chickpea linked to increased ADH levels in seeds at 10HAI in two temperatures:** early chickpea plant vigor is critical for plant productivity under terminal drought conditions (Sivasakthi et al., 2017). From the two principle chickpea types, Desi and Kabuli, vast field experience has shown that Desi is clearly superior in terms of multi-stress tolerance and yield performance (Purushothaman et al., 2014). We could in a former research discriminate both types at 10 HAI by a lower oxycaloric equivalent (Rq/RCO2) value due to differential carbon use and, thus, predict posteriori the known better yield stability of Desi (Gunasekaran et al. 2019). Here we show that Desi increased the level of ADH at 10 HAI during germination (significant at 23°C and 28°C), while this was not seen in Kabuli. The reached level of ADH was higher at 23°C than at 28°C. **B.5 Pronounced SHAM-effects on ADH levels at 24HAI that show interaction with sucrose**: during the germination of carrot seeds, the most pronounced effect of SHAM-treatment on ADH levels was observed at 24 HAI. At that time point, SHAM stimulated ADH level compared to levels observed at 12 HAI and 30 HAI. This happened independent of the presence of sucrose (3%). However, under both conditions, 5 mM SHAM showed a stronger stimulating effect on ADH levels (significant) at 24 HAI than 10 mM SHAM. But the level of SHAM-enhanced ADH was higher at both the tested concentrations of SHAM when sucrose was not present. To the contrary, at both time points 12 HAI and 30 HAI, a higher ADH level in the 0% sucrose controls was associated to higher concentration of SHAM 10mM versus 5mM SHAM. At 3% sucrose, ADH activity was at any time point higher at 5mM SHAM than at 10mM SHAM. Together, these observations point to the importance of differential AOX activity-regulation for optimized germination during all three time points independently on the presence or absence of exogenous sucrose. **C.1 SHAM pulses ≤12HAI impact germination efficiency and interact with sucrose effects**: in control seeds, short pulses of SHAM (10mM) until 12 HAI enhanced germination efficiency and were more effective than pulses until 6 HAI. However, prolonged SHAM treatment of 72 HAI suppressed early germination. In contrast, at 3% exogenous sucrose, early germination efficiency was reduced against 0% sucrose controls (confirming Figure 1.A.1) and SHAM pulses until 6 HAI and 12 HAI led to complete suppression of early germination. However, from 48 HAI onwards to 72 HAI, continuous SHAM treatment in the presence of 3% sucrose increased germination, while under 0% sugar continuous SHAM suppressed germination also at 72 HAI. Collectively, these results show that plastic AOX regulation was critical for the timing of germination in controls and under conditions of sucrose supplementation. **C.2 10HAI and 30-40HAI are critical times for sucrose-SHAM interaction during carrot germination**: 10 h of previous water imbibition reduced the strong negative effects of a combination of exogenous sucrose (3%) and SHAM (5 mM) on germination efficiency that were observed at only 2 h of previous water imbibition (significant). Also during the phase of initiated root emergence at 30 HOI (hours of imbibition) a transfer from water to media supplemented with sucrose and SHAM suppressed germination (significant). Water imbibition until 40 h before transfer to sucrose- and SHAM-containing media relieved and even supported germination when monitored at 30 HAI and at 48 HAI (significant). However, this increase in germination efficiency seemed to be restricted from 72 HAI onwards (significant). **D. Sucrose and SHAM can improve the effect of AMF on early germination:** in Figure 1.D.1, it can be seen that carrot seeds, which were treated with native endophytes (isolated from cv. Kuroda) tended to improve early germination at 48HAI in both the cultivars (not seen for EN3 in cv. Early Nantes). Exogenous sucrose had differential effects depending on endophyte and cultivar (Figure 1.D.2), but in no case could sucrose enhance early germination rates compared to the respective endophyte-treated control (see also Supplementary Table S2). However, SHAM treatment (Figure 1.D.2) reduced early germination against endophyte-treated controls in all cases (see also Table S2). In a separate trial, two AMF strains (M1 and M2) from the species *Rhizophagus* were tested and acted negatively on germination in cv. Kuroda, but positively in slowly germinating seeds of cv. Early Nantes (Figure 1.D.1). Nevertheless, the effect of M1 on early germination could be improved in cv. Kuroda by 0.5% sucrose and 3% sucrose (Figure 1.D.2). However, this was not seen for M2. In the better germinating cv. Kuroda, the lower concentration of 5 mM SHAM (Figure 1.D.2) improved the effect of both mycorrhiza species on early germination. In later germinating seeds of cv. Early Nantes, 0.5% sucrose improved the already positive effect on germination (Figure 1.D.1) of *Rhizophagus* strain M1 (Figure 1.D.2). In this cultivar, SHAM decreased the germination rate to the level of the untreated control (Table S2). E.1. ***On-farm* organic vs conventional seed discrimination and organic breeding by help of quick germination tests, commercial sugar and SHAM treatment:** seeds from 6 of 7 winter wheat cultivars originating from organic agricultural management could be discriminated at 15 HAI through better germination against conventionally produced seeds when germinated in 5% sugar solution. In water, seeds of only 4 cultivars showed better germination for organic seeds. When conventionally produced material was compared, seeds of cultivar 1 showed poor germination. This was much more pronounced, when tested in 5% sugar solution instead of water. Seeds of cultivar 2 demonstrated highest germination rates among all tested cultivars (see Figure S4). This was observed for seeds originating from both agricultural conditions, although higher germination in 5% sugar solution indicated the presence of microbiota (Figure 1.E.1). In contrast to all other cultivars, seeds of cultivar 2 did not differ in germination rates for organic vs conventional production under SHAM treatment when compared to the water control (see also Figure S4). This signals already low levels of *AOX* at 15 HAI for this cultivar no matter from which agricultural management system seeds originated. Overall, these observations indicate interplay between plant genotype, sugar and AOX activity that impacts differential germination capacities between organic and conventional seeds. E.2. **Identification of disease tolerant pea seeds by germination tests under SHAM-discrimination (T - tolerant reference; S – susceptible reference):** Pea lines with differential degrees of root rot disease susceptibility could be ranked by employing SHAM-inhibition. The most tolerant line (T) showed the lowest degree of SHAM-related inhibition of germination monitored at 27 HAI. This indicates the reasonability of germination tests under SHAM discrimination for selection of seed vigor and plant robustness. E.3. ***On-farm* simple seed germination improvement of stored conventional and organic wheat seeds by a 2h initial pulse with commercial sugar** (cultivar 1 in 1.E.1): this figure demonstrates the general potential of improving early germination through a short pulse of sugar and its validity across species (here winter wheat, see for carrot Figures 1.A.2 and S2), agricultural management practices and also related to the aging of seeds.

**In summary**, we found that **(a)** during *Arabidopsis thaliana* seed germination *ADH* transcript levels increased 12 h after seed stratification in water followed by a decline and that increase in ADH transcript levels was in general accompanied by increased *AOX1a* transcript accumulation (Figure 1.B.2) **(b)** in agreement with **(a)**, germinating carrot seeds displayed a higher level of biochemically determined ADH at 12 HAI than at 24 HAI. In the presence of 3% sucrose, this level was enhanced (Figures 1.A.3 and 1.B.3) **(c)** short pulses of sucrose of 2h at water imbibition enhanced early germination in seeds of two different species, *viz*., carrot and wheat (Figures 1.A.2, S2, 1.E.3). In carrot, we showed that the effectiveness of such early sugar pulse was dependent on sucrose concentration. A short pulse could be substituted by a longer pulse at a lower concentration of sucrose (Figure 1.A.2) **(d)** to the contrary, SHAM treatment until 6 HAI and 12 HAI suppressed germination in the presence of 3% sucrose. However, it favored early germination in the absence of sucrose (Figure 1.C.1).

**(e)** Three carrot native bacterial endophytes were used for carrot seed inoculation on two cultivars and showed a tendency to improve germination (Figure 1.D.1). However, a positive effect was dependent on cultivar-endophyte interaction. SHAM treatment reduced early germination percentage of endophyte-treated seeds against the respective endophyte-treated controls. This was observed in all cases though to a different degree (Figure 1.D.2 and table S2) **(f)** sucrose had differential impacts on endophyte-mediated effects on germination and was dependent on cultivar and endophytes. However, in no case endophytes improved germination of sucrose-treated seeds to higher levels than the endophyte-treated controls without sucrose (Figure 1D.2 and table S2) **(g)** In a good germinating carrot cultivar, the two selected *Rhizophagus* strains acted both negatively on early germination, while in a later germinating carrot cultivar, both *Rhizophagus* strains acted positively (Figure 1.D.1 and table S2). Sucrose could improve *Rhizophagus* effects on early germination to higher levels than the AMF-treated controls in both the cultivars. However, this was dependent on cultivar-strain interaction. In the presence of sucrose, strain M1 improved germination of both cultivars compared to M1-treated control seeds (Figure 1.D.2). **(h)** at lower concentrations of SHAM (5 mM), early germination could be improved to higher levels as compared to the AMF-treated controls (Figure 1.D.2), but this was observed only in the better germinating cultivar, which had not shown positive AMF effects against non AMF-treated controls (Figure 1.D.1 and table S2)

In **Figure 2**, we present a simplified scheme that summarizes our conclusions based on wet-lab experiments, state-of-the-art knowledge and our hypothetical inferences related to the dynamic metabolic interplay between sucrose, aerobic fermentation, cyt respiration, AOX regulation/alternative respiration, and microbiota on cell reprogram functioning. In this scheme, we separate AOX as a macromolecule (gene/protein) from the functional pathway, the alternative respiration, to highlight the outstanding position of AOX as the key and only enzyme of a pathway that, if present in an organism, was recognized to provide a central metabolism-coordinating function for efficient survival (Mohanapriya et al., 2019, Arnholdt-Schmitt et al., 2021 In Press, Costa et al., 2021; preprint). We consider that under development- and/or environment-induced conditions of rapid sucrose increase, the Cyt pathway is stimulated via enhanced glycolysis, pyruvate production and increased TCA cycling in a way that the respiration chain can get overloaded by electrons followed by enhanced ROS/RNS levels and, on the other hand, restricted due to rapidly consumed oxygen and/or yet low numbers of functional mitochondria in relation to available oxygen during germination. In turn, aerobic alcoholic and lactic fermentation are stimulated (see points a), b) and c), and Costa et al., 2021; preprint). At the same time, AOX is activated (see point d) and in Mohanapriya et al., 2019, Costa et al., 2021; preprint) mainly through AOX gene sequence-dependent pyruvate regulation and ROS/RNS.

**Figure 2:**
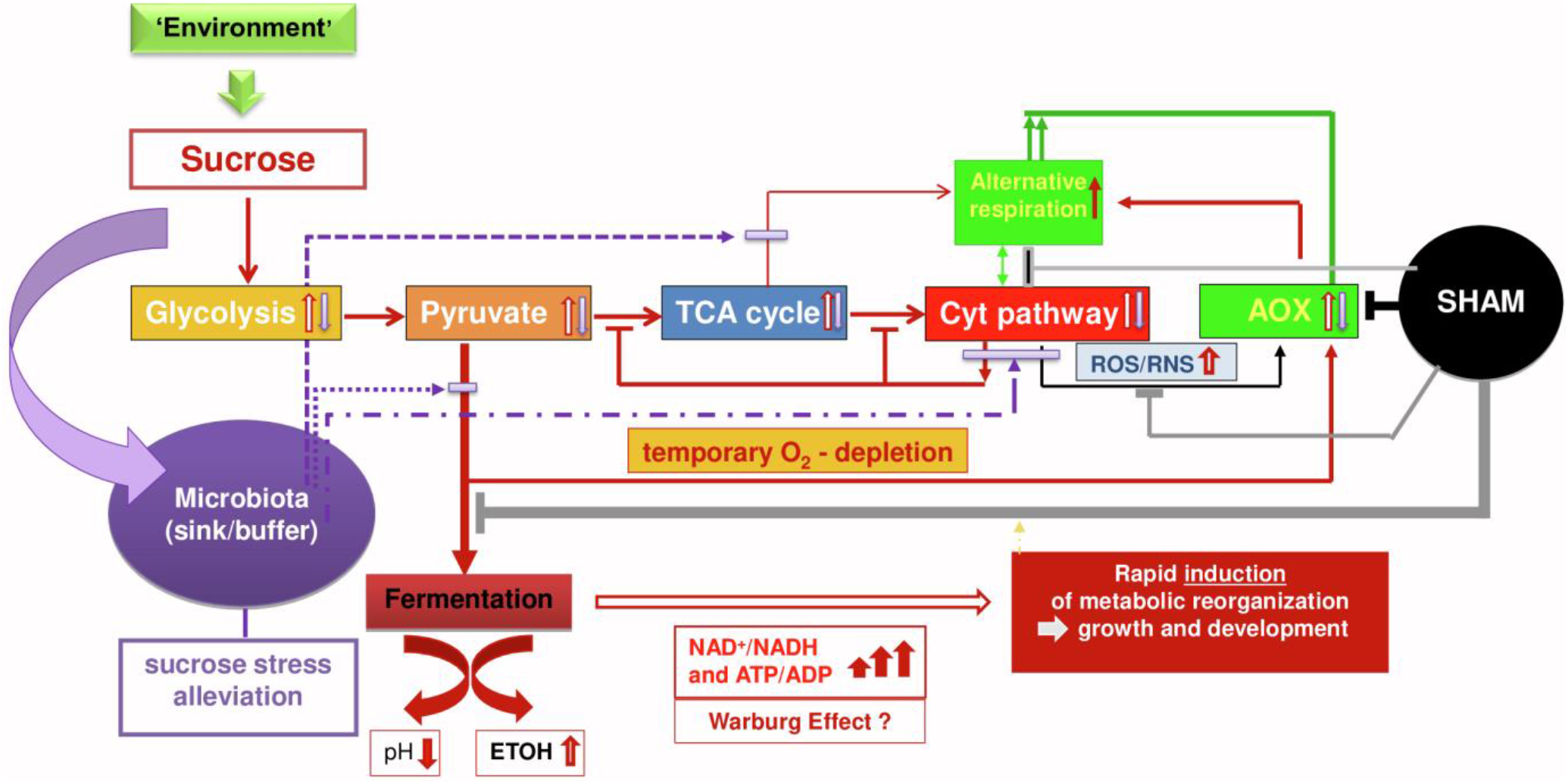
A simplified scheme on hypothesis and conceptualization for working out metabolic principles on dynamic cell reprogram-functioning (details explained in text)

Depending on stress level and the amount of sucrose and duration of a situation of high sugar-level, anaerobic glycolysis can reach high turnover during cell reprogramming and a level of high ATP production even corresponding to the Warburg effect. This latter hypothesis is supported by a parallel research on auxin-induced callus growth (Costa et al., 2021; preprint) where we observed a rapid and high increase in *ADH1* transcripts of 1777% and a parallel increase in *LDH* transcripts of 346%. Warburg effects are increasingly recognized also in human systems (Melkonian and Schury, (2020), Kutschera et al., 2020) as being part of normal physiology. However, in plants they are studied still more in relation to photosynthesis (Kutschera et al., 2020) and anaerobic conditions are best explored in relation to anaerobic conditions under flooding and was related to anaerobic tolerance in rice (Narsai et al., 2017). It was shown that, AOX plays beneficial role under low oxygen and especially during re-oxygenation (Jayawardhane et al., 2020).

Under increased sucrose, fermentation can escape feedback down-regulation by the help of enhanced alternative respiration, since AOX-transferred electrons enable continuation of TCA cycling for metabolic re-organization though with a relatively less energy efficiency. Thus, fermentation and AOX are complementing each other in order to maintain metabolic and energetic homeostasis thereby avoiding inefficient situations when the respiration chain is overloaded in relation to oxygen availability. As soon as oxidative stress gets sufficiently diminished at equilibrated oxygen availability in the cyt pathway, AOX will be down-regulated and normal respiration will reach priority again for driving growth and development. Fermentation and AOX will again be regulated in adaptation to sucrose- and cyt respiration-transmitted conditions embedded in adaptive hormonal crosstalk and overall complex cellular and apoplastic network signaling. Thus, rapid down-regulation indicates efficient adaptation of cyt respiration, a dynamic trait appropriate to mark seed vigor (Mohanapriya et al., 2019).

Sucrose can improve early germination of *Rhizophagus*-treated seeds (see point g) while non-AMF-treated seeds respond upon sucrose typically with a delay in germination (see Figure 1.A.1). This suggests that AMF can alleviate or buffer negative effects of sucrose on germination to relevant degrees by providing an additional sink. This was not indicated for the three tested endophytes (f). Also, early germination of endophyte-treated seeds was reduced at 48 HAI by continuously present SHAM when compared to seed germination of the respective endophyte-treated controls (e). To the contrary, when seeds from the good germinating cultivar were inoculated with *Rhizophagus*, SHAM treatment (5 mM) could improve early germination to higher levels than observed in AMF-treated controls. This observation is in agreement with the palliating effects observed by *Scutellospora calospora* on negative SHAM effects on carrot germination by using the same cultivar (Mohanapriya et al., 2019). In an overall assessment, it is inferred that AMF treatment might improve early germination by alleviating stress by rapid sucrose excess through two mechanisms: providing an additional sink for sucrose and supplying an enhanced capacity and/or engagement of alternative respiration. *Rhizophagus* spores were shown to be a rich source for polymorphic AOX gene sequences (Campos et al., 2015). We believe that there could be operation of two separate mechanisms, since we observed differential effects on early germination of M1-treated seeds upon SHAM-treatment in the two selected cultivars (Figure 1.D). However, M1-treated seeds of both cultivars showed improvement in early germination when sucrose was provided (Figure 1.D). We tend to interpret that the isolated native carrot endophytes were already well integrated into the internal host cell habitat. Thus, their re-inoculation tended to influence early germination positively, but could not provide a striking new advantage or disadvantage when sucrose was enhanced or SHAM-treatment reduced the level of alternative respiration. However, we reported that endophytes modulate *AOX* transcripts in a species-, stress-, and development-dependent manner and that endophytes could have modified the effect of AMF inoculation on seed germination efficiency (Mohanapriya et al. 2019).

### Outlook

Our observations offer new perspectives for low-cost prediction of plant holobiont behavior from seeds and for providing simple and rapid *on-farm* support towards sustainable agriculture. We propose three tools for validation:

A) Seed selection by help of short germination tests under SHAM discrimination. This tool provides modalities to identify seeds with higher seed vigor, general adaptive plant robustness and superior internal seed quality related to the content of secondary metabolites (Figures 1.E.1, 1.E.2, S3 and S4)

B) Discrimination of organic versus conventionally produced seeds with the help of short duration germination tests in water solutions with 5% commercial sugar (Figure 1.E.1)

C) Germination improvement by 2 h pulses of commercial sugar (Figures 1.A.1, S1, 1.E.3)

Furthermore, we encourage developing novel tests for AMF functionality in germinating seeds in the presence of sucrose. This approach targets compatibility between selected plants and AMF strains to support plant holobiont plasticity.

Our results suggest that polymorphic AOX gene sequences of symbiotic partners can impact plant-AMF compatibility. Therefore, we want to accomplish wider screening of major *AOX* polymorphisms in species-specific target cells for evaluating plant performance (Abe et al., 2002, Arnholdt-Schmitt et al., 2006, Arnholdt-Schmitt, 2015, Nogales et al., 2016) and in AMF sources (Arnholdt-Schmitt, 2008; Vicente and Arnholdt-Schmitt, 2008, Campos et al., 2015). Such strategy needs to also include near neighboring polymorphisms in conserved functional sites that can discriminate differentially regulated *AOX1* and *AOX2* (Costa et al., 2009). This approach would include screening of compatible *AOX* polymorphisms from both partners in the proposed functional tests to identify best plant-AMF combinations.

We hypothesize that the observed integration of bacterial endophytes into host plants with similar sensitivity against SHAM effects might point to synchronized AOX regulation in plant holobionts. Into this derivation would fit that we observed the same tendency of inhibiting sucrose effects on endophyte-free and superficially sterilized seeds (Figure 1.A.1), which we noticed also for SE induction (unpublished). Vicente et al. (2015) highlighted a ‘provocative’ lack of interest in bacterial AOX. They anticipated that bacteria-harboring AOX could facilitate adaptation to extreme conditions, which could also be of interest when thinking on plant endophytes and AMF-associated bacteria (Pandit et al., manuscript under preparation).

This present perspective is complementing Mohanapriya et al. (2019) and Costa et al. (2021; preprint). Joining the central figures of these publications is thought as one teaching tool that can help explaining a straightforward way from fundamental interdisciplinary research to application that might support sustainable socio-economies in view of the diversity of emergent environmental changes.

## Author Contributions

BR performed lab analyses on carrot germination, endophyte isolation and inoculation trials related to Figures 1.A.1-3, 1.B.3, 4 and 5, 1.C.1 and 2, and 1.D. JHC coordinated transcriptome analyses supported by KTL. JHC, RS and CN discussed initially the approach of this manuscript with BAS. GM carried out work on Figure S1 and Table S1. SS was responsible for AMF inoculation trials under the head of AA, ESM performed pea studies for Figure E.2 under responsibility of BAS. Under supervision of KJG, ESM together with AK performed germination analyses of transgenic Arabidopsis, and AK carried out the ADH analyses on chickpea. BAS performed *on farm* analyses (Figures 1.E.1 and 1.E.3). CN was responsible for statistics and was in part supported by MO. BR and IV helped BAS in literature search. DS contributed with Figure S3. BAS initiated the scientific approach, coordinated overall research and discussion and wrote the manuscript. All co-authors commented research and manuscript during its development and agreed to manuscript submission. BR organized manuscript submission.

## Acknowledgments

RS, GM and BAS acknowledge support for academic cooperation and researchers mobility by the India-Portugal Bilateral Cooperation Program (2013-2015), funded by “Fundação para a Ciência e Tecnologia” (FCT), Portugal, and the Department of Science and Technology (DST), India. GM is grateful to UGC, India, for doctoral grant from BSR fellowship. KJG, MO and BAS acknowledge support by the India-Portugal Bilateral Cooperation Program ‘DST/INT/Portugal/P-03/2017’. MO Research is partially supported by National Funds through FCT, Fundação para a Ciência e a Tecnologia, projects UIDB/04674/2020 (CIMA). BAS wants to thank RS for enabling intensive external online supervision of BSR on the presented research and excellent collaboration and communication of BSR. BR and SS acknowledge the infrastructure and stay support provided by DBT-TDNBC-DEAKIN – Research Network Across continents for learning and innovation (DTD-RNA) for AMF related work at The Energy and Resources Institute, TERI, India. JHC is grateful to CNPq for the Researcher fellowship (CNPq grant 309795/2017-6). KTL is grateful to CNPq for the Doctoral fellowship. BAS is grateful to SRK for his support in facilitating coordination of the Indian FunCROP team. CN acknowledges the international scientific network BIOALI-CYTED, which contributed to establish FunCROP contacts. BAS wants to acknowledge especially the extraordinary engagement of CN for online collaboration with BSR on data evaluation and presentation and overall manuscript discussion. BAS appreciates collaboration of LIVESEED partners with seed material and information on this material and thanks for supporting ESM (European Horizon2020 project LIVESEED GRANT NO. 727230).

## Supplementary Tables

**S1: Table S1:** Effect of exogenous sucrose concentration on carrot SE callus induction

**S2: Table S2:** Microbiota effect on carrot seed germination at different sucrose and SHAM concentrations

## Supplementary figures

**S1:** Exogenous sucrose delayed callus emergence and was necessary for SE

**S2:** 2 h pulse with commercial sugar improved carrot germination efficiency monitored at 40 HAI and 50 HAI

**S3:** Effect of SHAM treatment on accumulation of soluble and wall bound phenolics (A) and flavonoids and lignin (B) in elicitor-treated hairy roots of *Daucus carota*. Values obtained in only elicitor-treated root was considered as 100% and results were expressed in terms of percentage of maximum. The terms E and NE in the x-axis legend denote -with and -without elicitor, respectively. * Soluble phenolics. Values are mean of three independent experiments ± SD.

**S4:** Rapid germination check of organic and conventional seeds from seven cultivars in water (control) or under SHAM (5 mM) treatment

**Supplementary file:** Materials and Methods

