## Supplementary Figure S1 and Supplementary Table S2 for "Adaptive reprogramming during early seed germination requires temporarily enhanced fermentation – a critical role for alternative oxidase (AOX) regulation that concerns also microbiota effectiveness"

|  | Callus induction efficiency  (in %) | |
| --- | --- | --- |
| Con. of Sucrose | SSS | EFS |
| 0% | 0±0 | 0±0 |
| 0.5% | 0±0 | 0±0 |
| 2% | 17±1.4 | 21.5±0.7 |
| 3% | 33.5±3.5* | 38±2.8* |
| 7% | 0±0 | 0±0 |
| p value | 0.025 | 0.0150 |

**Supplementary Figure 1:** Exogenous sucrose delayed callus emergence and was necessary for SE

c

c

c

b

a

b

b

b

b

a

c

bc

bc

a

a

c

bc

bc

b

a

c

c

b

b

a

d

d

c

b

a

d

d

c

b

a

c

a

b

b

ab

ab

b

ab

ab

No. induced calli

| Days after incubation | 0% Suc. | 0.5% Suc. | 2% Suc. | 3% Suc. | 7% Suc. |
| --- | --- | --- | --- | --- | --- |
| 1 | 1 | 0.666667 | 0.666667 | 0 | 0 |
| 2 | 2 | 1 | 0.666667 | 0 | 0 |
| 3 | 7 | 5 | 1.66667 | 0 | 0 |
| 4 | 19 | 19 | 13.3333 | 8.66667 | 0.666667 |
| 5 | 27.6667 | 25 | 20.3333 | 16 | 2 |
| 6 | 29 | 26.6667 | 20.6667 | 20.3333 | 2 |
| 7 | 31 | 28.6667 | 26.3333 | 23 | 5.33333 |
| 8 | 31 | 28.6667 | 27 | 23 | 8.33333 |
| 9 | 31 | 29 | 27.6667 | 24 | 10.3333 |
| 10 | 31 | 30.3333 | 29.6667 | 24.3333 | 16 |

The surface sterilized seeds used for SE callus inducton at different concentration of sucrose. Suc. – Sucrose; DAI – Days after inoculation; The bar represent the mean ± SD; Tukey test at the P>0.05.

**Supplementary Table S1:** Effect of exogenous sucrose concentration on carrot SE callus induction

|  | Callus induction efficiency  (in %) | |
| --- | --- | --- |
| Con. of Sucrose | Surface sterlized seeds | Endophyte free seeds |
| 0% | 0±0 | 0±0 |
| 0.5% | 0±0 | 0±0 |
| 2% | 17±1.4 | 21.5±0.7 |
| 3% | 33.5±3.5* | 38±2.8* |
| 7% | 0±0 | 0±0 |
| *p* value | 0.025 | 0.0150 |

SSS – Surface sterilized seeds; EFS – Endophytes free seeds, % - percentage; Values represent the mean ± SD; * indicates statistically significant (Student’s T – Test). The *p* value was compared between 2% and 3% sucrose on SE callus induction efficiency.
