## Supplementary Figure S2 for "Adaptive reprogramming during early seed germination requires temporarily enhanced fermentation – a critical role for alternative oxidase (AOX) regulation that concerns also microbiota effectiveness"

**Figure S2**: 2 h pulse with commercial sugar improved carrot germination efficiency monitored at 40 HAI and 50 HAI

| Hours after imbibition | No. Seeds germinated |
| --- | --- |
| H2O 40HAI | 4 |
| 2h sucrose 40HAI | 7.2 |
| H2O 50HAI | 9.2 |
| 2h sucrose 50HAI | 13.6 |

**

**
