## Supplementary Table S2 for "Adaptive reprogramming during early seed germination requires temporarily enhanced fermentation – a critical role for alternative oxidase (AOX) regulation that concerns also microbiota effectiveness"

| **Organisms**  **Treatments** | **EN1** | **EN2** | **EN3** | **AMF1** | **AMF2** |
| --- | --- | --- | --- | --- | --- |
| **cv. Kuroda** | | | | | |
| **0.5% sucrose** | **8.66 ± 1.66ab** | **10 ± 2.51bc** | **17 ± 0b** | **17.6 ± 1.6bc** | **7.3 ± 1.3ab** |
| **3% Sucrose** | **4.3 ± 0.33a** | **4.6 ± 1.2ab** | **17 ± 0.53b** | **17.3 ± 1.33abc** | **5.3 ± 1.33ab** |
| **5mM SHAM** | **7 ± 1.52ab** | **5 ± 1.15abc** | **10 ± 0a** | **22.3 ± 1.56c** | **13.6 ± 0.66bc** |
| **10mM SHAM** | **2.6 ± 1.2a** | **2 ± 0.5a** | **10 ± 0.5a** | **8.6 ± 1.33a** | **5 ± 1a** |
| **Treated Control** | **13.3 ± 0.8b** | **13 ± 2.3c** | **15.6 ± 0.6ab** | **13.3 ± 0.88ab** | **10.3 ± 2.33abc** |
| **Non-treated Control** | **11.6 ± 2.40b** | | | **20 ± 2.08bc** | |
| **cv. Early Nantes** | | | | | |
| **0.5% sucrose** | **4 ± 2ab** | **0.66 ± 0.66** | **0.33 ± 0.33 ab** | **2 ± 0.57** | **1.3 ± 0.66** |
| **3% Sucrose** | **0.66 ± 0.33a** | **0 ± 0a** | **0 ± 0** | **0.33 ± 0.33** | **0.33 ± 0.33** |
| **5mM SHAM** | **3.33 ± 1.76ab** | **0.33 ± 0.33** | **0.33 ± 0.33ab** | **0.66 ± 0.66** | **0 ± 0** |
| **10mM SHAM** | **0.66 ± 0.33a** | **0 ± 0a** | **0.33 ± 0.33** | **0 ± 0** | **0.66 ± 0.66** |
| **Treated Control** | **9.66 ± 0.8b** | **3 ± 1.5** | **0.66 ± 0.33 b** | **1.33 ± 0.88** | **1 ± 0** |
| **Non-treated Control** | **1.66 ± 0.33ab** | | | **0.66 ± 0.33** | |

**Table S2:** Microbiota effect on carrot seed germination at different sucrose and SHAM concentrations

Values represent the mean ± SD; Tukey test at the P>0.05.
