## Supplementary Figure S3 for "Adaptive reprogramming during early seed germination requires temporarily enhanced fermentation – a critical role for alternative oxidase (AOX) regulation that concerns also microbiota effectiveness"

**Figure S3**: Effect of SHAM treatment on accumulation of soluble and wall bound phenolics (A) and flavonoids and lignin (B) in elicitor-treated hairy roots of *Daucus carota*. Values obtained in only elicitor-treated root was considered as 100% and results were expressed in terms of percentage of maximum. The terms E and NE in the x-axis legend denote -with and -without elicitor, respectively. * Soluble phenolics. Values are mean of three independent experiments ± SD.

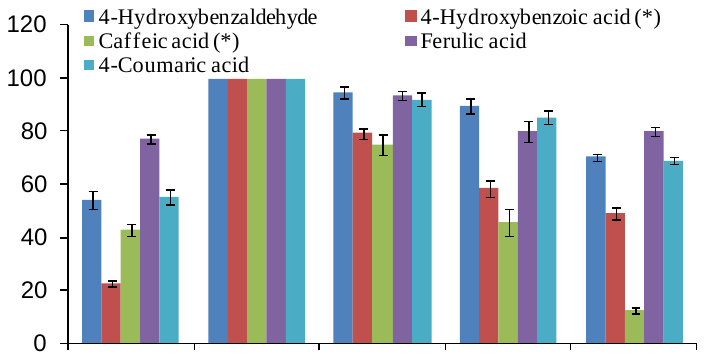

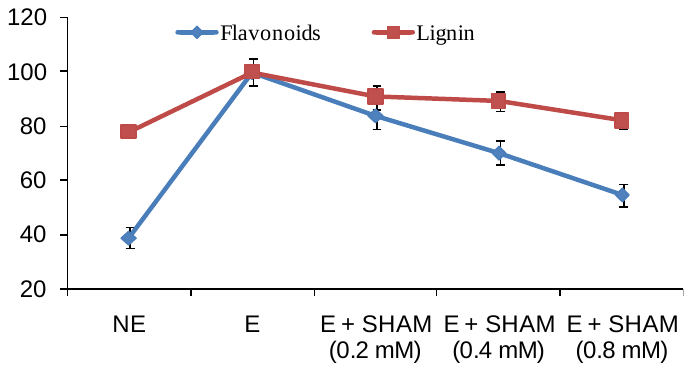

**% of maximum**

**A**

**B**
