## Supplementary Figure S4 for "Adaptive reprogramming during early seed germination requires temporarily enhanced fermentation – a critical role for alternative oxidase (AOX) regulation that concerns also microbiota effectiveness"

**Figure S4: Rapid germination check of organic and conventional seeds from seven cultivars in water (control) or under SHAM (5 mM) treatment**

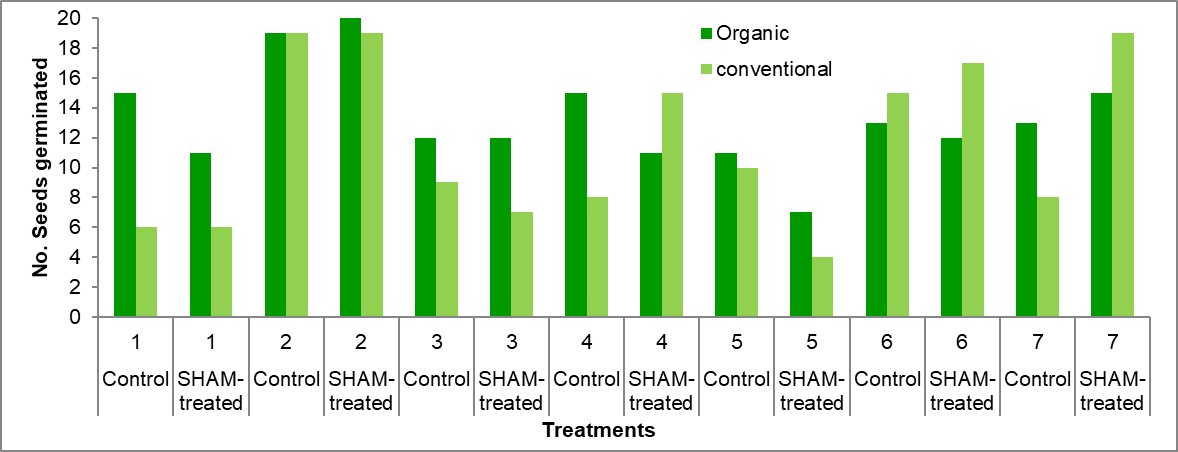
