## Supplementary material for "Adaptive reprogramming during early seed germination requires temporarily enhanced fermentation – a critical role for alternative oxidase (AOX) regulation that concerns also microbiota effectiveness": Materials and methods

**Material and Methods:**

**1. Carrot seed germination assays under sucrose and pulse treatments:**

Carrot seeds (varieties Kuroda and Early Nantes) were subjected to surface sterilization or native endophyte removal procedures as explained in Mohanapriya et al., (2019), see also under point 1.2 and 1.3. For germination, plates were incubated at 25˚C in dark for 40 hrs and then shifted to a culture room with 16/8 h (light/darkness) photoperiod provided by cool white fluorescent tubes. Germination was identified by radicle emergence and monitored at 24, 30, 48, 72, 96 and 120 HAI.

For sucrose treatments, different concentrations of sucrose were prepared (0.5%, 2%, 3%, 7%) using autoclaved deionized water. Sterile germination papers were placed onto petri dishes and 5 ml of respective sucrose solutions were added to the germination papers. 40 seeds were inoculated per petri dish and three replicates were maintained for each sucrose treatment.

For salicylhydroxamic acid (SHAM) treatment, SHAM was dissolved in dimethyl sulphoxide (DMSO) (Mohanapriya et al., (2019) made up with sterile distilled water. 5ml of 5 mM and 10 mM SHAM solutions were added onto sterile germination papers and 40 carrot seeds were inoculated. Three replicates were performed for each SHAM treatment. Deionized water served as the control. For SHAM-sucrose, sucrose + SHAM pulse experiments, seeds were inoculated in respective solutions of SHAM (5 mM, 10 mM) and sucrose (0.5%, 2%, 3%, 7%), sucrose + SHAM (5 mM SHAM without sucrose or 5 mM SHAM + 3% sucrose) and shifted to water at defined time points (2 HAI, 10 HAI, 30 HAI, 40 HAI) and vice versa.

**2. Bacterial endophyte and mycorrhiza-treatments of *D. carota* L. seeds:**

Seeds of two carrot varieties, Kuroda and Early Nantes, were pretreated to remove native endophytes by an established protocol (Mohanapriya et al., 2019). Briefly, stock solutions of (1 mg ml^-1^) tetracycline (antibiotic) and miconazole nitrate (antifungal) were prepared and 5 ml of each solution was transferred onto a Petri plate. 200 mg of carrot seeds were weighted and added into the plate containing antibiotic and antifungal solutions and kept under shaking (100 rpm) for 6 h. Thereafter, seeds were washed with sterile deionized water for thrice and kept under laminar air flow until they were completely dried. To confirm the effectiveness of the treatment, few seeds were cut into small pieces, placed on nutrient agar (Himedia, India) and incubated at 35^º^C overnight. Absence of microbial growth indicated that seeds were free from native endophytes.

For bacterial endophyte treatment, native bacterial endophytes were grown in autoclaved nutrient broth (HiMedia, India) at 35^º^C overnight. Then, cultures were diluted with sterile deionized water until the optical density (O.D.) reached 0.2 (2 x 10^8^ cells ml^-1^). The endophyte-free seeds were subsequently immersed in 20 ml of the respective bacterial cultures and kept under shaking at 100 rpm for 2 h. Seeds were dried under laminar air flow to remove excess of moisture.

For arbuscular mycorrhizal fungi (AMF) treatments, ninety days old root organ culture (ROC; stock culture) of two different *Rhizophagus species* were obtained from the Centre for Mycorrhizal Culture Collection (CMCC), TERI, India. Stock culture was harvested at 25°C in 100 ml of sodium citrate buffer (Doner and Becard 1991) using a shaker (Kuhner Shaker, Basel, Switzerland) for 60 min at 100 rpm. After deionization, buffer containing harvested spores (without roots) was sieved through a 325 British Standard (300 μm) Sieve (BSS) (Fritsch, Idar-Oberstein, Germany). Spores retained on the sieve mesh were washed with sterile distilled water (twice). Finally, the spores were collected in 20 ml of sterile distilled water. All steps were performed under aseptic conditions (Srivastava et al. 2016).

For experimental set-up, autoclaved germination papers (Glasil Scientific Industries, New Delhi, India) was placed onto Petri dishes and moisturized with 5 mM and 10mM SHAM. Sucrose solutions (0.5, 2, 3 and 7%) were used to moisturize the germination paper. 40 bacterial endophyte-treated seeds were placed on moisturized dishes using sterile forceps. In the case of the AMF treatments, 40 pretreated seeds were placed into the dishes and 5 spores were inoculated per seed separately for two different *Rhizophagus* species used in the study. All treatments had three replicates. Triplicates of controls were maintained by inoculating seeds with sterile distilled water.

**3. Induction of somatic embryogenesis and SHAM-treatment in *Daucus carota* L:**

Carrot seeds (varieties Kuroda) were subjected to surface sterilization or native endophyte removal procedures as explained in Mohanapriya et al., (2019), The dried seeds were collected into sterile screw-cap tubes and stored at room temperature. Surface sterilized or endophytes free seeds were inoculated on B_5_ solid medium supplemented with 0.5mg/L 2,4-D at different concentration of sucrose (0%, 0.5%, 2%, 3%, and 7%) according to previously established protocol (Mohanapriya et al., 2019). Plates were incubated in a culture room at 25˚C with 16/8 h (light/darkness) photoperiod. Triplicates of 40 seeds per plate were maintained for each treatment. The numbers of seeds response with induced callus were recorded 45 days after inoculation as well as seeds response at 1 to 10 day after inoculation were noted.

**4. Alcohol dehydrogenase assays:**

**4.1. ADH assay for carrot seeds:**

40 carrot seeds (cv. Kuroda) were cultured under SE-inducing conditions and collected at 0, 12, 24, 30, 48 and 72 HAI. They were pooled together and ground into powder by using liquid nitrogen in a sterile mortar and pestle. Fine powder was extracted using 1 mL of 1X phosphate saline buffer (pH 7.2) and centrifuged for 30 min at 10,000 rpm and 4 ^º^C. The supernatant obtained was used as the crude protein extract. The enzyme assay was performed according to the protocol given by Kagi and Vallee (1960). Briefly, the reaction mixture consisted of 1.3 ml 50 mM sodium phosphate buffer (pH 8.8), 0.1 ml 95% ethanol, 1.5 ml 5 mM ß-nicotinamide adenine dinucleotide (ß-NAD^+^) and 0.1 ml of crude protein extract. This mixture was immediately inversed several times for proper mixing and the 340 nm absorbance was recorded from 0 to 6 min. Alcohol dehydrogenase from *Saccharomyces cerevisiae* was used as the positive control. Active enzyme units (IU - international units) were calculated according to the following formula:

$\frac{(\Delta A340/min Test - \Delta A340/min Blank) (3.0) (df)}{(6.22) (0.1)}$

3.0 = Total volume (ml) of assay

df = Dilution factor

6.22 = millimolar extinction coefficient of reduced ß-nicotinamide adenine dinucleotide (ß-NAD^+^) at 340 nm

0.1 = Volume (ml) of crude extract used

In Figures 1.A.2 and 1.B.3 the calculated ADH units were multiplied by a factor (15 and 10 respectively) for better visualization of the relation between ADH and numbers of seeds with induced callus.

**4.2. ADH assay for Chickpea:**

Chickpea seeds of two varieties Pusa 362 Desi and Kabuli 1105 were surface sterilized with mercuric chloride, washed, imbibed in autoclaved distilled water and then kept on two different temperatures 23˚C and 28˚C for germination. For dry seed experiments the seeds were directly weighed kept on different temperature and used for the experiment. Approximately 100 mg of sample was weighed and crushed in liquid nitrogen and samples were used for ADH assay briefly seeds were weighed in triplicate and were crushed in liquid N2. The resulting powder was allowed to thaw at 4oC, after which it was centrifuged at 13,000 rpm for 15 minutes to remove cell debris, producing clarified supernatant for ethanol assay. Ethanol levels were determined using an enzymatic test kit according to the manufacturer’s instructions (K-ETOH kit; Megazyme International Ireland Ltd).

**5. Testing *on-farm* the influence of a 2 h-pulse of commercial sugar on carrot germination efficiency:**

Organic carrot seeds (cultivar Nantaise 2/Milan, Bingenheimer Saatgut (Germany)) provided by Steven P.C. Groot and Jan Kodde, Wageningen Plant Research, Wageningen University & Research, Wageningen, Netherlands, were used to perform an *on-farm* test on improving early germination efficiency by a 2h-pulse of commercial sugar (5% in water solution). A bulk of 40 seeds was germinated in Petri dishes at around 25°C room temperature (September 2019) on 4-layer kitchen paper and by using available water at a private site in Portugal. The trial was performed in five repetitions and macroscopically monitored at 40 HAI and 50 HAI.

**6. SHAM treatment of wildtype (WT) and transgene, *AOX1a*-silenced (AS) and *AOX1a* –overexpressed (OE) *Arabidopsis thaliana* seeds:**

For salicylhydroxamic acid (SHAM) treatment, SHAM was dissolved in dimethyl sulphoxide (DMSO) different concentrations of SHAM 0.5 mM, 1.5 mM and 2.5 mM was used. WT, *AOX1a*-silenced (AS) and *AOX1a* –overexpressed (OE) *Arabidopsis thaliana* seeds were surface sterilized with help of Triton X 100 and 70 % ethanol . Total three replicates were used for control and treatment. Afterwards seeds were placed on filter paper until it dr . After drying seeds were transferred on filter paper and sprayed with 2 ml of sterile water. Plates were sealed with micropore tape and placed at 4˚C for 48h for stratification. 2.5 ml of SHAM solution was added onto the plates , Control was treated with sterile water and phenotypic observations were made after 27, 48, and 72h.

**7. Expression analysis of *AtADH1+2*** **and *AtAOX* genes from transcriptomic data using RNA-seq:**

Gene expression was evaluated during germination of *Arabidopsis* using RNA-seq data collected from Sequence Read Archive (SRA) database of GenBank, NCBI, bioproject: PRJNA369750. The RNA-seq data were derived from *Arabidopsis* seeds according to the following time points in triplicates: Dry seeds (0 h), seeds after; 1 h of stratification (1 h S), 12 h of stratification (12 h S), 48 h of stratification (48 h S), followed by seed collected 1 hour into the light (1 h SL), 6 hours into the light (6 h SL), 12 hours into the light (12 h SL), 24 hours into the light (24 h SL) and 48 hours into the light (48 h SL). More experimental details about RNA-seq samples can be found in Narsai et al. (2017).

The expression analyses was carried out for *AtADH1, 2* and *AtAOX* (*AOX1a, 1b, 1c, 1d* and *2*) genes, which specific regions (3’ end) of each cDNA were used to map reads in RNA-seq data according to Saraiva et al. (2016). To ensure a more accurate count of mapped reads an additional analysis was included using the Magic-Blast (Boratyn et al., 2019). The resulting data (number of mapped reads) were normalized using the RPKM (Reads Per Kilobase of transcript per Million of mapped reads) method (Mortazavi et al. 2008). Normalized data were obtained by applying the following equation: RPKM = (number of mapped reads X 109) / (number of sequences in each database X number of nucleotides of each gene).

**8. *On-farm* check of wheat seeds from organic vs conventional agriculture:**

Seeds of seven winter wheat cultivars were tested in a rapid *on-farm* check to discriminate seeds that originated from parallel production in the same region under organic and conventional agriculture. Originally, all seeds came from conventional production in 2017, but had been divided and finally harvested in 2018 from both types of agricultural management. The material was made available by Peter Mikó, Agricultural Institute of the Centre for Agricultural Research Hungarian Academy of Sciences, Hungary. Cultivars were anonymously numbered from 1 to 7, considering that these trials had been preliminary *on-farm* checking that should not be taken as a definite cultivar characterization produced in a specific region.

A bulk of twenty seeds was germinated in petri dishes on 4-layer kitchen paper by using *on-farm* available water on a site in Portugal at around 25°C room temperature in June 2019. In parallel, germination was performed in water, a water solution of 5% commercial sugar and in a water solution that was substituted after 2 h by 5 mM SHAM. Germination was macroscopically observed as root emergence at 15 HAI.

In order to check whether germination of stored seeds could be improved by a short pulse of sugar (5% water solution), independently on their origin from organic or conventional production, a bulk of twenty seeds from cultivar 1 was germinated in December 2020 in 5 Petri dishes (repeats) on 4-layer kitchen paper by using *on-farm* available water on a site in Portugal at around 20°C room temperature. Germination was macroscopically observed as root emergence at 15 HAI for conventional seeds and at 18 HAI for organic seeds, because of the reduced germination efficiency of organics seeds from this cultivar despite storage under identical conditions.

**9. Pea line discrimination by help of SHAM:**

Seeds of twenty *Pisum sativum* accessions with known resistance/susceptibility performance against pea root rot disease had been used for SHAM discrimination studies. The material had been made available through FiBL, Schwitzerland within the frame of the European LIVESEED project (European Horizon2020 project LIVESEED GRANT NO. 727230). However, these trials were an agreed project extension beyond direct FiBL collaboration. Original seed sample identification numbers had been changed to 1-18 plus a tolerant and susceptible accession considering also that these analyses are preliminary studies to develop the methodology, but should not be taken for granted characterization of the material.

10 seeds per Petri dish were germinated in 3 replicates on 4-layer kitchen paper with double distilled water at 25ºC and humidity of 58% in a climate chamber (Fitoclima D1200PLH, Aralab, Portugal). At 10 HAI, seeds were transferred to a 10 mM SHAM double distilled water solution. Germination had been macroscopically monitored as root emergence.

**10. Statistical analyses**

Normality and homogeneity of variances from the analyzed variables were tested with Shapiro-Wilk test and Bartlett or Levene tests, respectively. If data were parametric, Student’s t test (two populations) or 1- or 2-way ANOVA were used. ANOVA tests were followed by Tukey´s post hoc tests when a significant effect of the tested factor was detected. If data were not parametric, Wilcoxon (two populations) or Kruskal-Wallis tests were performed. Statistical packages used were InfoStat 2018I and R v. 4.0.2.

We highlight that we interpret our data as ‘real’ observations under the employed conditions involving only small samples, which certainly provide insights that cannot get relevance or not relevance by using significance calculation. Nevertheless, we applied significance calculations at usual *p*-values for biological research as an aid to appropriately focus our insights. Readers are encouraged to making themselves familiar with the current paradigm change related to the usage of statistical significance (Hirschauer and Becker, 2020 and references herein).
